# Robust IgM responses following vaccination are associated with prevention of *Mycobacterium tuberculosis* infection in macaques

**DOI:** 10.1101/2021.05.06.442979

**Authors:** Edward B. Irvine, Anthony O’Neil, Patricia A. Darrah, Sally Shin, Alok Choudhary, Wenjun Li, William Honnen, Smriti Mehra, Deepak Kaushal, Hannah Priyadarshini Gideon, JoAnne L. Flynn, Mario Roederer, Robert A. Seder, Abraham Pinter, Sarah Fortune, Galit Alter

**Affiliations:** Ragon Institute of MGH, MIT and Harvard; Cambridge, MA, USA; Harvard T.H. Chan School of Public Health; Boston, MA, USA; Vaccine Research Center, National Institute of Allergy and Infectious Diseases (NIAID), National Institutes of Health; Bethesda, MD, USA; Public Health Research Institute, New Jersey Medical School, Rutgers, The State University of New Jersey; Newark, NJ, USA; Department of Medicine, University of Massachusetts Medical School; Worcester, MA, USA; Division of Microbiology, Tulane National Primate Research Center; Covington, Louisiana, USA; Southwest National Primate Research Center, Texas Biomedical Research Institute; San Antonio, TX 78245, USA; Department of Microbiology and Molecular Genetics and Center for Vaccine Research, University of Pittsburgh School of Medicine; Pittsburgh, PA, USA

## Abstract

Development of an effective tuberculosis (TB) vaccine has suffered from an incomplete understanding of the correlates of protection against *Mycobacterium tuberculosis* (*Mtb*). However, recent work has shown that compared to standard intradermal Bacille Calmette-Guerin (BCG) vaccination, intravenous (IV) BCG vaccination provides nearly complete protection against TB in rhesus macaques. While studies have focused on cellular immunity in this setting, the antibody response elicited by IV BCG vaccination remains incompletely defined. Using an agnostic antibody profiling approach, here we show that IV BCG drives superior antibody responses in the plasma and the bronchoalveolar lavage fluid (BAL). While IV BCG immunization resulted in the broad expansion of antibody titers and effector functions, surprisingly, IgM titers were among the strongest markers of reduced bacterial burden in the plasma and BAL of BCG immunized animals. Moreover, IgM immunity was also enriched among animals receiving protective vaccination with an attenuated *Mtb* strain. Finally, a LAM-specific IgM monoclonal antibody reduced *Mtb* survival *in vitro*. Collectively, these data highlight the potential importance of IgM responses as a marker and as a functional mediator of protection against TB.

## INTRODUCTION

*Mycobacterium tuberculosis* (*Mtb*), the causative agent of tuberculosis (TB), was responsible for the death of an estimated 1.4 million individuals in 2019 (*1*). While TB is curable, the intensive antibiotic regimen coupled with the rise in antibiotic resistance has underscored the need for an efficacious vaccine to help mitigate the global TB epidemic. Bacille Calmette-Guérin (BCG), first introduced in 1921, is the current standard for TB vaccination (*2*). While BCG is effective at preventing severe forms of TB in young children, BCG is poorly and variably efficacious in preventing pulmonary TB in adults (*3*). Consequently, novel vaccines and vaccination strategies are urgently needed.

TB vaccine development has suffered from a lack of understanding of the determinants of immunity against *Mtb* infection. CD4+ T cell knock-out studies in mice (*4*, *5*), CD4+ depletion and simian immunodeficiency virus infection studies in non-human primates (NHPs) (*6*–*8*), as well as human immunodeficiency virus infected human cohort studies (*9*), have all found increased rates of TB disease progression in the setting of low CD4+ T counts, pointing to a critical role for CD4+ T cells in controlling *Mtb* infection. However, efforts to generate vaccines that primarily leverage T cell immunity to drive protection against TB have been met with limited success.

The first large-scale TB vaccine clinical trial since BCG was the T_H_1-directed MVA-85A vaccine phase 2b trial (*10*). Despite the lack of efficacy of MVA-85A vaccination, a post-hoc correlates analysis of the study found Ag85A-specific IgG responses to be linked with reduced risk of TB disease (*11*), identifying humoral immunity as an unexpected negative correlate of TB disease risk. More recently, the M72/AS01_E_ phase 2b TB vaccine trial in adults reported a 50% reduction in rates of progression to active TB (ATB) (*12*). While robust T cell immunity was observed following M72-vaccination, strong anti-M72-specific humoral immunity was also observed, as all vaccinees in the M72/AS01_E_ group remained seropositive 36 months following vaccination (*12*, *13*). Together, these human vaccination studies point to the potential importance of both cellular and humoral immunity in vaccine-mediated protection against TB.

In addition to the promising efficacy signals that have begun to emerge in human TB vaccine studies (*13*, *14*), intravenous (IV) BCG vaccination of non-human primates (NHPs) has been shown to provide robust protection against *Mtb* infection (*15*–*20*). Remarkably, IV BCG vaccination resulted in a 100,000-fold reduction in lung bacterial burden compared with standard intradermal BCG vaccination, with six out of ten macaques demonstrating no detectable level of *Mtb* infection (*20*). In contrast, high-dose intradermal BCG and aerosol BCG vaccination regimens resulted in lung bacterial burden levels similar to those observed with standard intradermal BCG vaccination (*20*). IV BCG vaccinated rhesus macaques exhibited increased antigen-responsive CD4 and CD8 T cells systemically, and locally in the lung compared to the other vaccination groups (*20*). Concomitant with enhanced T cell responses, IV BCG vaccinated animals elicited stronger whole-cell lysate reactive antibody responses in the plasma and bronchoalveolar lavage (BAL) fluid (*20*). Yet, while broad differences in antibody responses across BCG vaccination groups were observed in this study, their antigenic targets, functional and anti-microbial activity, and relationship with *Mtb* burden were not defined.

Here we leveraged Systems Serology to investigate the antigen-specific humoral immune responses that uniquely evolve following IV BCG vaccination in rhesus macaques (*21*). We demonstrate that compared to standard intradermal BCG vaccination, high dose intradermal BCG vaccination and aerosol BCG vaccination, IV BCG vaccination elicits superior antigen-specific humoral immunity in the periphery, and was the only regimen to induce a robust, lung-compartmentalized antibody response capable of restricting *Mtb* growth *in vitro*. While IgG, IgA, and several Fc-receptor binding antibody subpopulations expanded selectively in the lungs of IV immunized animals, antigen-specific IgM titers were strongly associated with reduced bacterial burden in the BAL and plasma of animals immunized with IV BCG. IgM titers were also found to be a marker of protective immunity in rhesus macaques mucosally vaccinated with an attenuated *Mtb* strain (*Mtb-ΔsigH*) – an orthogonal vaccination strategy also shown to protect rhesus macaques from lethal *Mtb* challenge (*22*). Finally, we show that a LAM-specific IgM monoclonal antibody reduced *Mtb* survival *in vitro*, suggesting that vaccine-induced IgM responses are plausible contributors to vaccine-induced protection against *Mtb*.

## RESULTS

### IV BCG immunization drives higher and more durable plasma antigen-specific antibody titers

Following IV BCG immunization, there was a marked increase in *Mtb* whole-cell lysate reactive IgG and IgA titers compared to BCG administration by other routes (*20*). However, in this first study, the antibody responses elicited by the different BCG vaccination strategies to distinct *Mtb* antigen targets were not assessed. Thus, we sought to determine whether particular antigen-specific antibody populations are differentially induced by different BCG vaccination strategies. Antibody levels to a panel of *Mtb* antigens were compared using a custom, multiplexed Luminex assay (*23*). The antigen panel included: purified protein derivative (PPD) – a heterogenous collection of *Mtb* proteins (*24*), lipoarabinomannan (LAM) – a critical cell wall glycolipid (*25*), HspX – a stress induced intracellular protein (*26*), as well as PstS1 and Apa – both cell membrane associated glycoproteins linked to host cell invasion (*27*, *28*). Of note, each of these antigens are expressed by both BCG and *Mtb*. Plasma samples collected pre-vaccination, week 8 post-vaccination, time of *Mtb* challenge (week 24), and post-infection (week 28) were analyzed, and fold change in antibody titer over pre-vaccination levels was calculated for each macaque at each timepoint.

Following immunization and prior to infection, antigen-specific IgG1 responses were detected in macaques across all vaccine arms. There were weak responses to all antigens in animals receiving standard ID BCG (Fig 1A). Conversely, those that received IV BCG vaccination displayed the largest increase in plasma IgG1 titers to nearly all tested antigens following vaccination. More specifically, PPD, LAM, PstS1, and Apa IgG1 titers in the IV BCG group were each significantly higher than those in the standard ID BCG group both at week 8 post-vaccination, and at the time of *Mtb* challenge (week 24 post-vaccination) (Fig 1A). The additional vaccination groups – high dose intradermal (ID_high_), aerosol (AE), and aerosol/intradermal (AE/ID) – trended towards higher IgG1 levels compared to the standard ID BCG group, though the differences were only significant for the ID_high_ Apa-specific response (Fig 1A).

**Figure 1:**
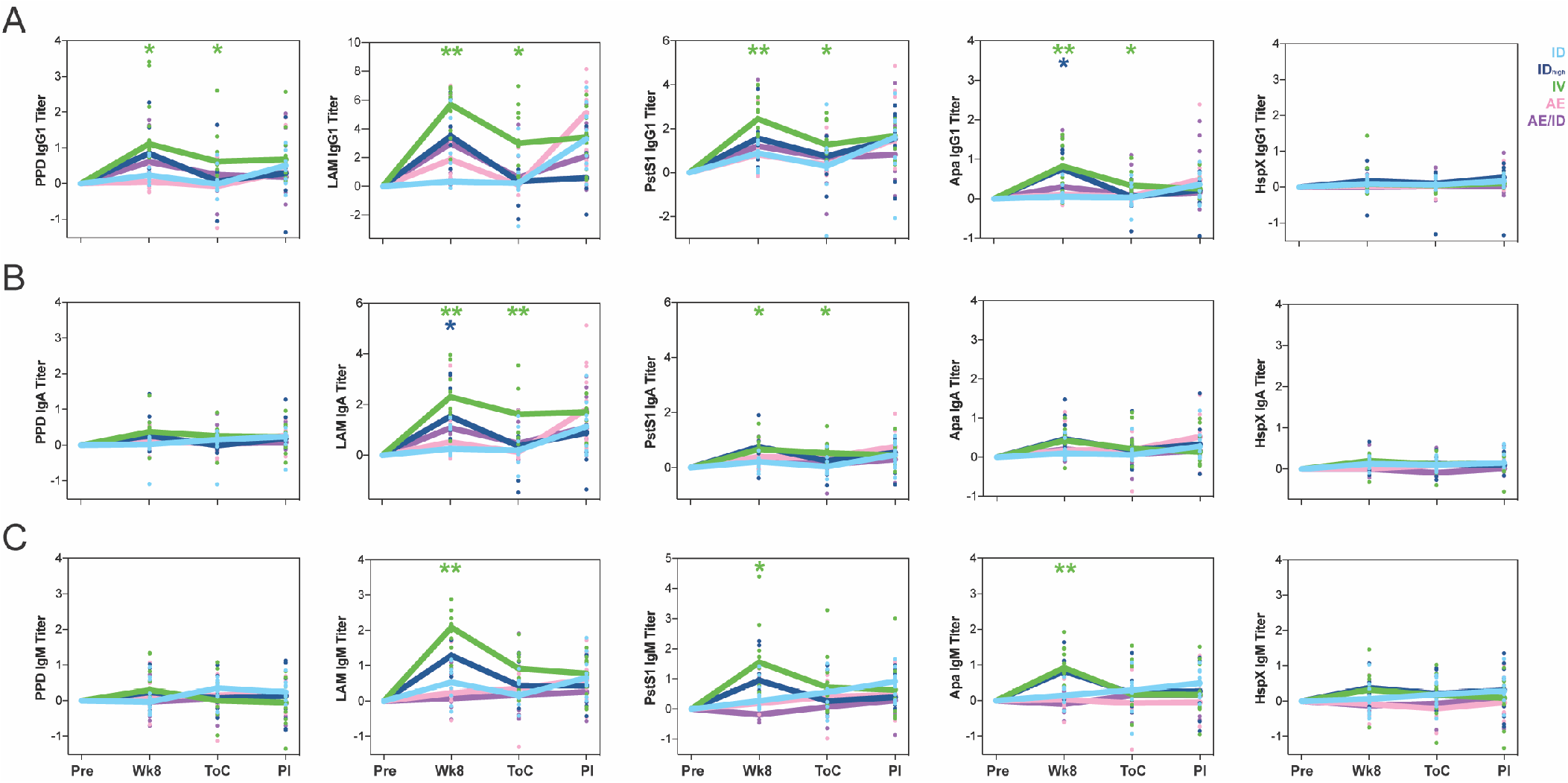
IV BCG immunized primates exhibit higher and more durable plasma antibody titers. Fold change in (**A**) IgG1, (**B**) IgA, and (**C**) IgM titers present in the plasma of each rhesus macaque following BCG vaccination were determined via Luminex. Fold changes were calculated as fold change in Luminex median fluorescence intensity (MFI) over the pre-vaccination level for each primate. A base-2 log scale is used for the y-axis. Timepoints: pre-vaccination (Pre), week 8 post-BCG vaccination (Wk8), time of challenge at week 24 post-BCG vaccination (ToC), post-infection at week 28 post-BCG vaccination (PI). Groups: standard intradermal BCG (light blue), high intradermal BCG (dark blue), intravenous BCG (green), aerosol BCG (pink), aerosol + intradermal BCG (purple). Each dot represents a single animal at the respective timepoint. The lines represent group medians over time. Kruskal-Wallis with Dunn’s multiple-comparison tests were performed on the fold change values at each timepoint, comparing each vaccination group to the standard intradermal BCG group. Adjusted p-values are as follows: *, p < 0.05; **, p < 0.01; ***, p < 0.001; ****, p < 0.0001.

Antigen-specific IgA responses were also observed following vaccination across each of the experimental BCG vaccination groups, though the fold increases in IgA titer were not as prominent as those for IgG1. IV BCG vaccinated macaques elicited significantly higher IgA titers to LAM and PstS1 at week 8 post-vaccination and at the time of *Mtb* challenge compared to the standard ID BCG group, which did not generate a detectable increase in antigen-specific IgA titers to any of the antigens following vaccination (Fig 1B). Animals in the ID_high_ group also elicited a significant increase in LAM IgA titers at week 8 post-vaccination (Fig 1B). However, minimal vaccine-induced plasma IgA responses to the additional antigens were detected in the other groups (Fig 1B).

Finally, antigen-specific IgM responses were detected in multiple experimental BCG vaccination groups, with IV and ID_high_ vaccinated macaques mounting the strongest peripheral IgM responses to BCG vaccination. IV BCG vaccinated animals exhibited significantly higher LAM-, PstS1-, and Apa-specific IgM titers at week 8 following vaccination compared to the standard ID BCG group, while responses in the ID_high_ group were not significantly different (Fig 1C). LAM IgM titers also trended higher at the time of challenge in the IV group, though the difference was no longer significant at this timepoint (Fig 1C).

Together, these data highlight peripheral differences in the antibody response to specific antigens induced by distinct BCG vaccination strategies. IV BCG immunized animals generated particularly high antigen-specific antibody levels in the plasma, with protein- and LAM-specific antibody responses persisting exclusively in IV immunized animals 24 weeks following vaccination, to the time of *Mtb* challenge. Further, standard ID BCG vaccination generated a weaker antigen-specific antibody response than the experimental vaccination regimens tested – each of which delivered a larger dose of BCG in addition to changing route – suggesting that altering both route and dose may result in enhanced peripheral humoral immune responses to BCG vaccination.

### IV BCG vaccination uniquely elicits a robust lung-compartmentalized antibody response

We next aimed to profile the antigen-specific humoral immune response at the site of infection using bronchoalveolar lavage fluid (BAL) collected from each macaque pre-vaccination, week 4 post-vaccination, and week 16 post-vaccination – the final timepoint the BAL procedure was performed prior to *Mtb* challenge. IV BCG vaccination uniquely elicited a robust antibody response in the airways following vaccination (Fig 2A – C). Specifically, IV BCG vaccinated animals mounted IgG1, IgA, and IgM responses in the BAL that were significantly higher than the standard ID BCG group at week 4 across all mycobacterial antigens tested (Fig 2A – C). The magnitude of the responses was particularly striking, with over 100-fold increases in antibody levels observed for some IV BCG vaccinated macaques (Fig 2A). Most antibody responses elicited were transient, with only statistically significant levels of LAM-, PstS1-, and Apa-specific antibodies detected in the BAL 16 weeks following vaccination. A small number of macaques in the ID_high_, AE, and AE/ID groups additionally generated detectable antibody titers in the BAL following BCG vaccination (Fig 2A – C). However, these responses were limited to one or two macaques in each group, and were substantially lower in magnitude than responses generated in IV BCG immunized animals (Fig 2A – C). These data indicate that IV BCG vaccination alone induced strong, lung-compartmentalized, antigen-specific humoral immune responses. These antibodies contracted, but persisted at detectable levels in IV immunized animals for at least 4 months following immunization.

**Figure 2:**
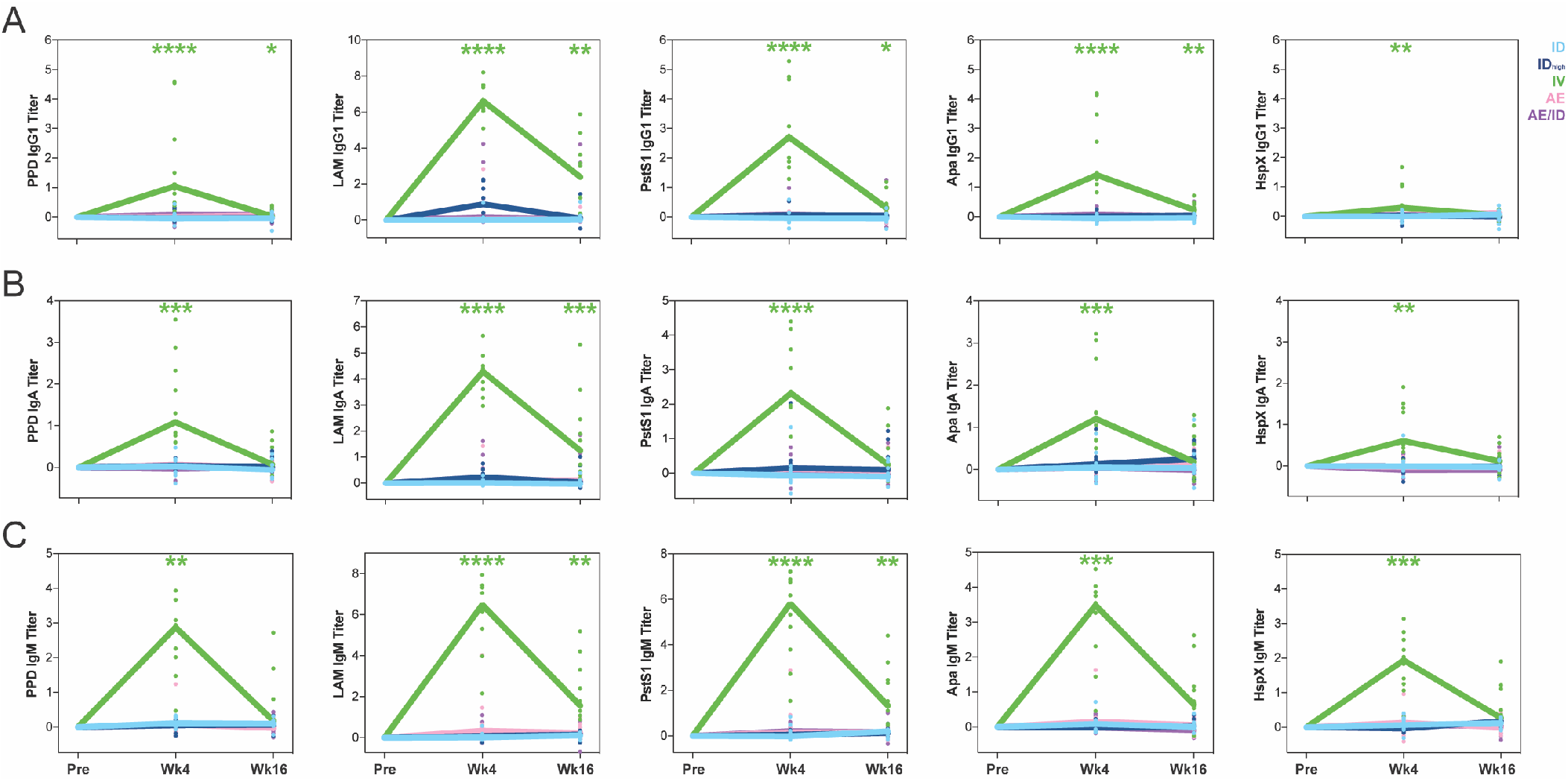
IV BCG vaccination uniquely elicits a robust lung-compartmentalized antibody response. Fold change in (**A**) IgG1, (**B**) IgA, and (**C**) IgM titers present in the BAL of each rhesus macaque following BCG vaccination were determined via Luminex. Fold changes were calculated as fold change in Luminex MFI over the pre-vaccination level for each primate. A base-2 log scale is used for the y-axis. Timepoints: pre-vaccination (Pre), week 4 post-BCG vaccination (Wk4), week 16 post-BCG vaccination (Wk16). Groups: standard intradermal BCG (light blue), high intradermal BCG (dark blue), intravenous BCG (green), aerosol BCG (pink), aerosol + intradermal BCG (purple). Each dot represents a single animal at the respective timepoint. The lines represent group medians over time. Kruskal-Wallis with Dunn’s multiple-comparison tests were performed on the fold change values at each timepoint, comparing each vaccination group to the standard intradermal BCG group. Adjusted p-values are as follows: *, p < 0.05; **, p < 0.01; ***, p < 0.001; ****, p < 0.0001.

### Antibodies from IV BCG vaccinated macaques mediate superior innate immune activation

Beyond their ability to bind and recognize pathogens or pathogen-infected cells, antibodies are able to deploy the anti-microbial activity of the innate immune system via Fc:Fc-receptor engagement to control a wide range of microbes (*29*). Further, Fcγ receptor (FcγR) signaling is necessary for the optimal survival and bacterial containment of *Mtb* in mice (*30*). Thus, we next measured the FcγR binding and functional capacity of plasma- and BAL-derived antibodies elicited in each BCG-vaccinated macaque to determine whether certain antibody FcγR binding profiles and/or antibody effector functions selectively tracked with distinct BCG vaccination strategies.

In the plasma, the IV BCG group displayed a trend towards higher levels of FcγR binding antibodies than the standard ID BCG group across nearly all antigens measured, including significantly increased PPD-, PstS1-, and Apa-specific FcγR2A and FcγR3A binding antibodies 8 weeks post-vaccination (Fig 3A). Further, although FcγR binding antibodies waned by the time of *Mtb* challenge across all vaccination groups, IV BCG vaccinated macaques maintained significantly higher levels of PPD- and PstS1-specific FcγR2A and FcγR3A binding antibodies close to the time of challenge, suggesting durable antibody functionality in this group (Fig 3A). To examine plasma antigen-specific antibody functionality, antibody-dependent phagocytosis by monocytes and neutrophils, as well as NK cell degranulation assays were performed. Each of these measurements were captured for LAM-specific antibodies, as each BCG vaccination regimen elicited detectable LAM-specific antibody titers in the plasma (Fig 1A – C). Antibodies from IV BCG vaccinated macaques induced the most potent antibody-dependent neutrophil phagocytosis, which was moderately, yet significantly higher than that observed in the standard ID BCG group at week 8 post-vaccination (Fig 3B). In contrast, limited differences were observed in LAM-specific antibody-dependent monocyte phagocytosis and antibody-dependent NK cell degranulation – a surrogate for antibody-dependent cellular cytotoxicity (ADCC) (*31*) – across the vaccine groups (Fig 3B).

**Figure 3:**
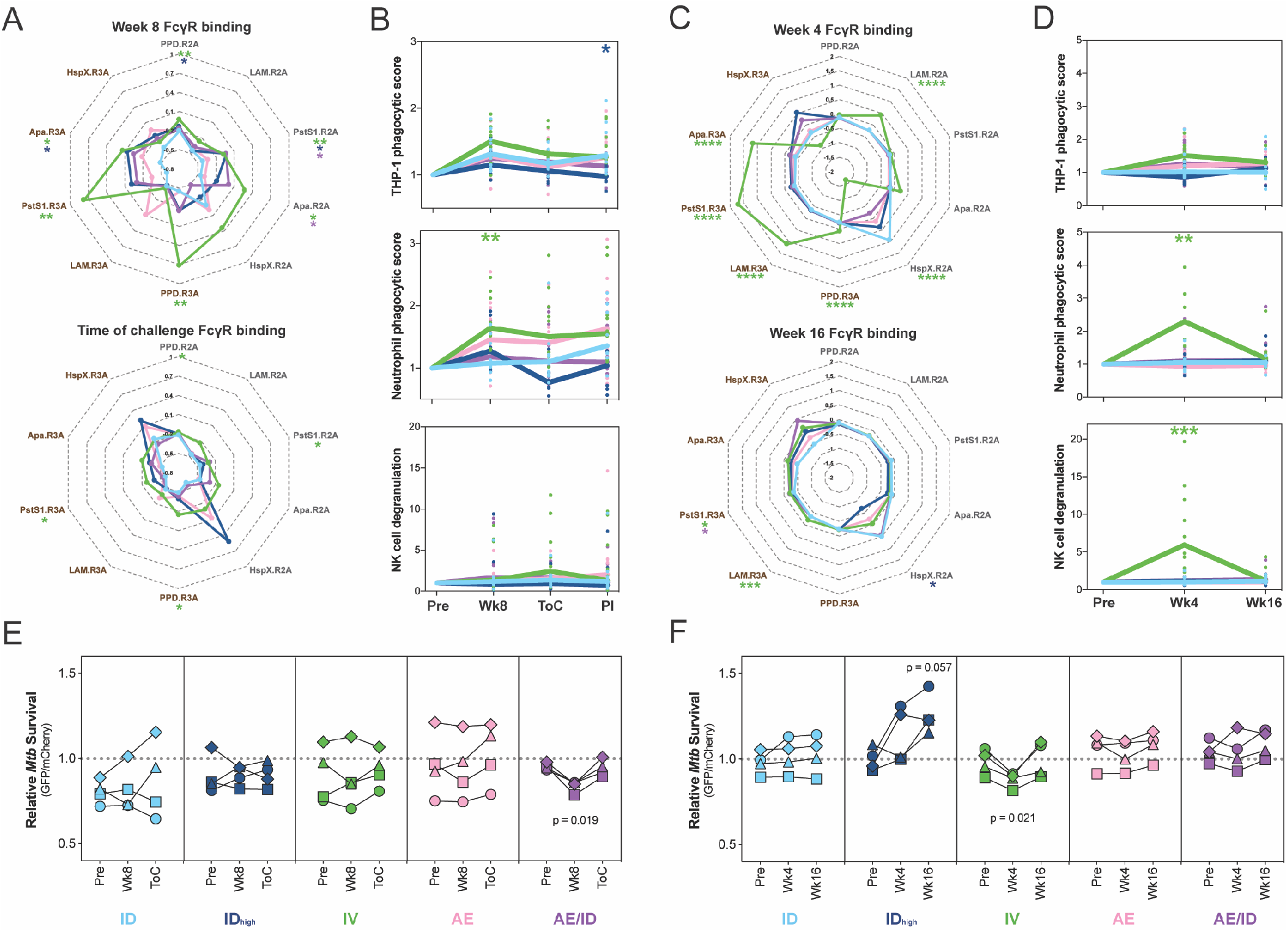
Antibodies from IV BCG vaccinated primates drive innate immune activation. (**A and C**) Radar plots of fold change in (**A**) plasma and (**C**) BAL antibody FcγR binding activity of each group post-BCG vaccination. Fold changes were calculated as fold change in Luminex MFI over the pre-vaccination level for each primate. Median z-score of each group is plotted for a given feature. (**B and D**) Fold change in (**B**) plasma and (**D**) BAL antibody-dependent cellular phagocytosis by THP-1 cells (top), antibody-dependent neutrophil phagocytosis by primary human neutrophils (middle), antibody-dependent primary human NK cell degranulation determined by % of cells CD107a positive (bottom). Fold changes were calculated as fold change over the pre-vaccination level for each primate. Each dot represents a single animal at the respective timepoint. The lines represent group medians over time. (**E and F**) *in vitro* macrophage *Mtb* survival assay using pooled (**E**) plasma and (**F**) BAL from each vaccination group at each timepoint. y-axis shows live (GFP) / total (mCherry) *Mtb* burden in human monocyte-derived macrophages. Lower on the y-axis indicates increased intracellular *Mtb* killing in macrophages. Each set of connected dots indicates the activity of pools from different timepoints run across the same healthy human macrophage donor; 4 donors were run in total. Plasma timepoints: pre-vaccination (Pre), week 8 post-BCG vaccination (Wk8), and time of challenge at week 24 post-BCG vaccination (ToC). BAL timepoints: pre-vaccination (Pre), week 4 post-BCG vaccination (Wk4), week 16 post-BCG vaccination (Wk16). Groups: standard intradermal BCG (light blue), high intradermal BCG (dark blue), intravenous BCG (green), aerosol BCG (pink), aerosol + intradermal BCG (purple). (**A – D**) Kruskal-Wallis with Dunn’s multiple-comparison test was performed on the fold change values at each timepoint, comparing each vaccination group to the standard intradermal BCG group. (**E and F**) Repeated measures ANOVA with Dunnett’s multiple comparisons test was performed for each vaccination group, comparing pre-vaccination restrictive activity with that of each post-vaccination timepoint. Adjusted p-values are as follows: *, p < 0.05; **, p < 0.01; ***, p < 0.001; ****, p < 0.0001.

In line with the elevated antibody levels observed in the BAL of IV BCG vaccinated macaques (Fig 2A – C), antibodies in the IV group demonstrated the highest levels of FcγR binding (Fig 3C). IV BCG immunized animals generated significantly higher levels of LAM-specific FcγR2A binding antibodies at week 4 post-vaccination (Fig 3C). In addition, antigen-specific FcγR3A binding levels in the BAL were particularly robust in the IV vaccinated group, with IV macaques displaying significantly higher levels of FcγR3A binding antibodies to PPD, LAM, PstS1, and Apa 4 weeks following vaccination (Fig 3C). LAM-, and PstS1-specific FcγR3A binding antibody levels remained significantly higher at week 16 post-vaccination (Fig 3C). Furthermore, BAL-derived antibodies in the IV group demonstrated superior LAM-specific functional activity. Specifically, BAL-derived antibodies from IV BCG vaccinated animals exhibited a trend towards stronger antibody-dependent monocyte phagocytosis activity (Fig 3D). More strikingly, the IV BCG group demonstrated significantly higher antibody-dependent neutrophil phagocytosis and NK cell degranulation activity week 4 vaccination, with little functionality observed in the other vaccination groups (Fig 3D). This activity returned to baseline in a majority of animals by week 16 post-vaccination (Fig 3D).

Previous data have linked an enrichment of FcγR3A binding and NK cell activating antibodies in the setting of LTBI, to enhanced intracellular *Mtb* killing in macrophages (*32*). Therefore, given the expansion of both of these humoral features particularly in the BAL of the IV immunized group 4 weeks post-vaccination, we next examined the anti-microbial activity of antibodies from each vaccination group in this context. Human monocyte-derived macrophages were infected with a live/dead reporter strain of *Mtb* (*33*), followed by the addition of pooled plasma or BAL from each BCG vaccination group. Plasma from the IV group did not drive significant *Mtb* restriction across either of the timepoints (Fig 3E). Conversely, the week 4 IV BCG BAL pool did drive moderate, yet significant intracellular *Mtb* restriction, whereas the ID_high_ BCG BAL pool tended to enhance infection. These patterns were consistently observed across all tested macrophage donors (Fig 3F).

Taken together, these data highlight the induction of highly functional antibodies following IV BCG immunization in rhesus macaques. Further, the increases selectively observed in FcγR3A binding, NK cell degranulation, and intracellular *Mtb* killing in the BAL were particularly salient given recent associations reported between both FcγR3A binding, as well as NK cell activity, and improved *Mtb* control (*32*, *34*).

### Antigen-specific IgM titers in the plasma and BAL negatively correlate with *Mtb* burden

A spectrum of bacterial burden was observed in the lungs of rhesus macaques across the BCG vaccinated groups at the time of necropsy (*20*). Thus, despite IV immunization clearly affording optimal bacterial control following *Mtb* challenge, we next aimed to define whether any antibody features exhibited a robust relationship with lung *Mtb* burden.

In the plasma, 5 antibody measurements were significantly negatively associated with *Mtb* burden after multiple hypothesis testing correction (Fig 4A) (*35*). Surprisingly, each of the features identified were antigen-specific IgM titers at week 8 post-vaccination or at the time of challenge (Fig 4A and B), revealing an unexpected significant relationship between plasma antigen-specific IgM titers and improved outcome following *Mtb* challenge. In contrast, while higher antibody titers have historically been associated with elevated antigenic burden and enhanced *Mtb* disease, antibody levels and features were not identified that tracked positively with *Mtb* burden at either significance level (Fig 4A).

**Figure 4:**
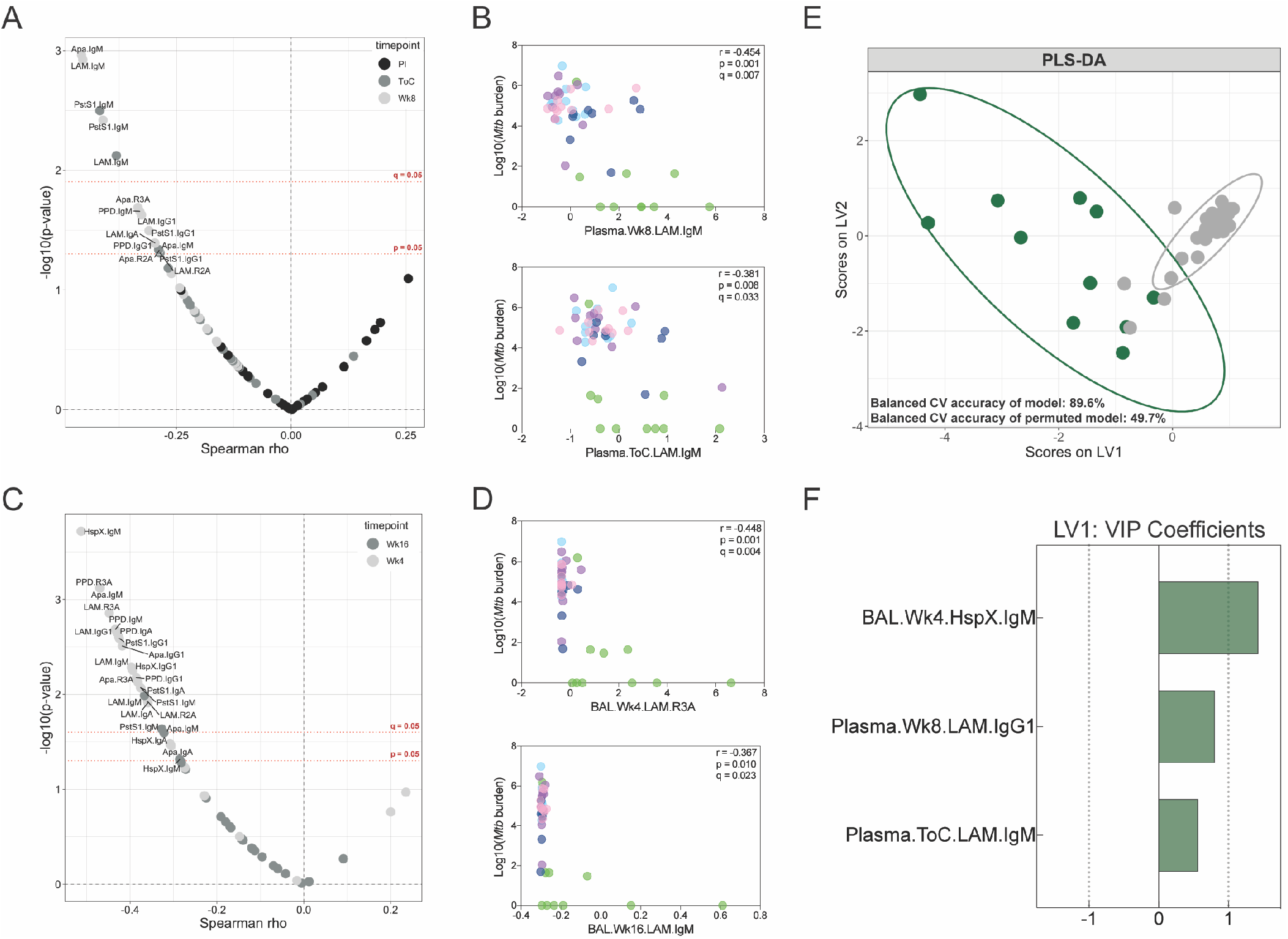
Numerous BCG-induced antibody features are associated with reduced *Mtb* burden. (**A and C**) Spearman correlations between base-10 log(*Mtb* burden) at necropsy and each (**A**) plasma and (**C**) BAL antibody measurement post-vaccination were computed. The x-axis indicates the spearman rho value associated with a given antibody feature. The y-axis indicates the negative base-10 log of the p-value associated with a given antibody feature. Antibody features are colored by their timepoint. (**A**) Plasma colors: week 8 post-BCG vaccination (light grey), time of challenge at week 24 post-BCG vaccination (dark grey), post-infection at week 28 post-BCG vaccination (black). (**C**) BAL colors: week 4 post-BCG vaccination (light grey), week 16 post-BCG vaccination (dark grey). (**B and D**) Spearman correlations between base-10 log(*Mtb* burden) at necropsy and select (**B**) plasma and (**D**) BAL antibody measurements. Points are colored by vaccination group: standard intradermal BCG (light blue), high intradermal BCG (dark blue), intravenous BCG (green), aerosol BCG (pink), aerosol + intradermal BCG (purple). Fold change antibody measurements were subjected to a z-score transformation prior to correlation analyses. Adjusted p-values (q-values) computed by the Benjamini-Hochberg procedure. (**E and F**) PLS-DA model fit using the antibody features selected by LASSO regularization. (**E**) Graph of the first two latent variables (LVs) of the model. Protected primates (green) had an *Mtb* burden < 1000 at necropsy. Susceptible primates (grey) had an *Mtb* burden > 1000 at necropsy. Ellipses show 95% confidence intervals. Balanced cross-validation (CV) accuracy of the model and permuted model are indicated. Accuracy of model is significantly higher than that of the permuted model (Mann-Whitney U test, p < 2.2e^−16^). (**F**) Variable importance in the projection (VIP) coefficients on LV1 for each model feature, indicating the extent to which each feature contributes to separation along LV1.

In the BAL, 18 antibody features were significantly negatively associated with *Mtb* burden after multiple hypothesis testing correction (Fig 4C) (*35*). Several antigen-specific IgG1, IgA, IgM, and FcγR binding measurements in the BAL at 4 or 16 weeks post-vaccination were negatively correlated with *Mtb* levels at the time of necropsy (Fig 4C and D). The majority of these features associated with reduced *Mtb* bacterial burden included antibody features present week 4 post-vaccination, the exception being LAM and PstS1 IgM titers at week 16 (Fig 4C and D). Again, none of the BAL antibody features measured had a significant positive correlation with *Mtb* burden at necropsy at either significance level (Fig 4C).

Collectively, the particularly low *Mtb* burden present in IV immunized animals indicate that the relationship observed between select humoral features and bacterial burden track with vaccination route, and thus may not represent independent correlates of protection. Nevertheless, these analyses point to the vaccine-induced humoral immune features which track most closely with improved microbial control in this vaccination cohort. Notably, IgM responses alone tracked with reduced *Mtb* burden close to the time of challenge across both compartments, potentially representing direct mechanistic correlates of immunity, or markers of a unique functional humoral immune response in these animals.

### Antibody profiles accurately distinguish protected and susceptible BCG-vaccinated macaques

Given that many antibody titer and functional measurements were highly correlated, even across compartments, we next sought to determine whether a minimal set of antibody features could be defined that collectively tracked with *Mtb* control. Thus, macaques with a lung *Mtb* burden at necropsy below 1000 were categorized as protected (total n = 11; 9 IV BCG, 1 ID_high_, 1 AE/ID BCG), and those with an *Mtb* burden greater than or equal to 1000 were categorized as susceptible (total n = 37). Next, least absolute shrinkage and selection operator (LASSO) regularization was implemented on the standardized antibody data, removing variables unrelated to the outcome, as well as reducing the number of highly correlated features (*36*). Partial least squares discriminant analysis (PLS-DA) was then performed to visualize and quantify group separation (*37*, *38*).

Robust separation was observed between protected and susceptible macaques on the basis of humoral profile (Fig 4E). The model distinguished protected from susceptible animals with a balanced cross-validation accuracy of 89.6% (Fig 4E). Remarkably, only 3 features were required to achieve this high level of predictive accuracy: BAL HspX-specific IgM at week 4, plasma LAM-specific IgG1 at week 8, and plasma LAM-specific IgM at the time of challenge. Each of these features contributed to separation along latent variable 1 (LV1) (Fig 4F). The selection of these three variables across distinct timepoints suggests that substantive humoral differences were present between protected and susceptible BCG-vaccinated macaques beginning in the lung in week 4, extending out to the time of challenge in the plasma. Further, this analysis demonstrates that protected and susceptible BCG-vaccinated macaques can be accurately resolved by simply using antibody titer measurements.

### Protective vaccination via attenuated *Mtb* is associated with increased plasma IgM titers

While our analyses identified humoral features associated with reduced *Mtb* burden in BCG immunized animals, because vaccination route was so closely linked to protection in this cohort, the generalizability of these findings was unclear. Thus, we next queried whether similar humoral features were associated with *Mtb* control in an independent *Mtb* vaccination study in NHPs. Previous work demonstrated that AE vaccination with an attenuated *Mtb* strain (*Mtb-ΔsigH*) provided superior protection compared to AE BCG vaccination in rhesus macaques (*22*). Thus, antibody profiling was performed on the plasma of *Mtb-ΔsigH* or AE BCG vaccinated animals.

Using antibody titer measurements alone, *Mtb-ΔsigH* and BCG vaccination groups could be clearly separated using a principal component analysis (PCA) (Fig 5A). Analysis of the PCA loadings plot revealed that antigen-specific IgM responses primarily drove separation between the two groups, with antigen-specific IgM responses enriched among protected *Mtb-ΔsigH* vaccinated macaques (Fig 5B). Similarly, univariate analyses indicated that *Mtb-ΔsigH* vaccinated macaques elicited significantly higher LAM-specific IgM titers week 7 post-vaccination, as well as a trend towards increased Apa- and HspX-specific IgM titers (Fig 5C). In contrast, minimal differences in antigen-specific IgG1 and IgA titers were noted between the *Mtb-ΔsigH* and BCG groups (Fig S4). Finally, antibody responses to the Ebola virus negative control antigen were not detected in either group as expected regardless of isotype (Figs 5C and S4).

**Figure 5:**
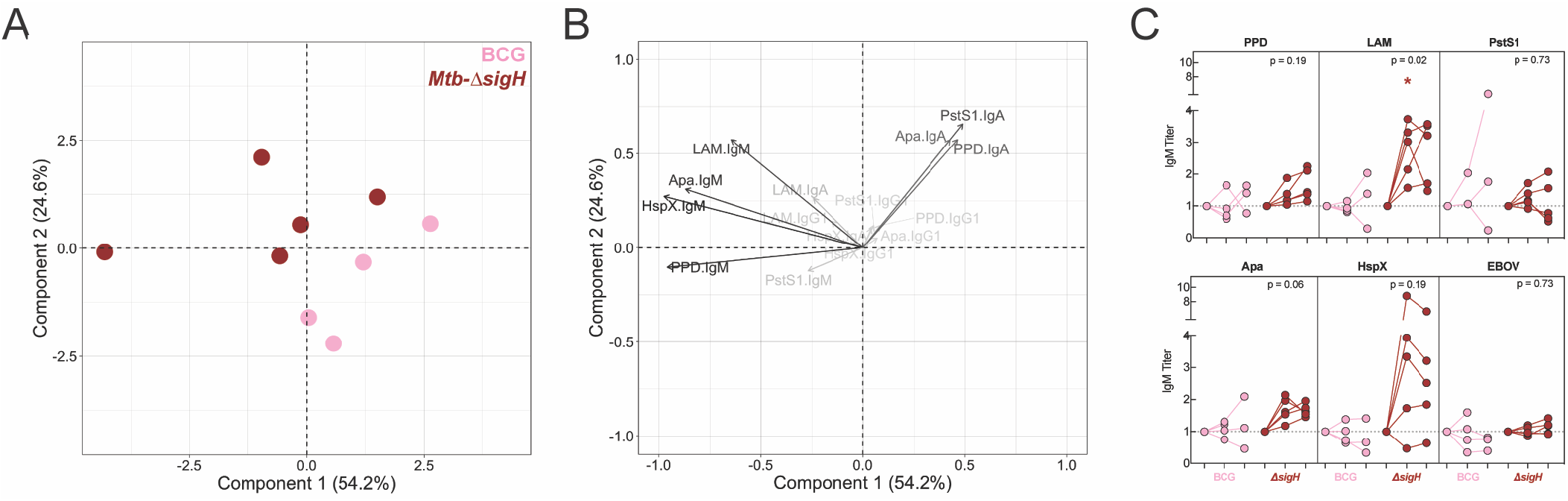
Protective vaccination with attenuated *Mtb* (*Mtb-ΔsigH*) is associated with increased plasma IgM titers. (**A and B**) Principal component analysis using fold change IgG1, IgA, and IgM titers measured at week 7 post-vaccination in an attenuated *Mtb* rhesus macaque vaccination cohort. Fold changes were calculated as fold change in Luminex MFI over the pre-vaccination level for each primate. Fold change antibody measurements were subjected to a z-score transformation prior to principal component analysis. (**A**) Score plot. Aerosol *Mtb-ΔsigH* vaccination group (red), aerosol BCG vaccination group (pink). (**B**) Loading plot. Relative contribution of variables to the components are indicated by a color gradient. Light grey variables contribute least, black variables contribute most. (**C**) Fold change in IgM titer present in the plasma of each rhesus macaque following vaccination determined via Luminex. Fold changes were calculated as fold change in Luminex MFI over the pre-vaccination level for each primate. Each dot represents a single animal at the respective timepoint. Timepoints: pre-vaccination (left), week 7 post-vaccination (middle), week 15 post-vaccination at necropsy (right). *Mtb* challenge was performed week 8 post-vaccination. Mann-Whitney U test was performed on the fold change values at each timepoint, comparing aerosol *Mtb-ΔsigH* to the aerosol BCG group. p-values are as follows: *, p < 0.05; **, p < 0.01; ***, p < 0.001; ****, p < 0.0001.

Thus, although the sample size from this cohort is small, increased plasma antigen-specific IgM titers tracked with reduced *Mtb* disease. A result similar to that observed in BCG immunized animals (Fig 4A), potentially hinting at a common association between antigen-specific IgM and vaccine-induced *Mtb* control.

### Superior *in vitro* anti-microbial effect of LAM-specific IgM

Data from both the BCG route and the *Mtb-ΔsigH* immunization study pointed to an unexpected association of antigen-specific IgM titers with improved vaccine-induced *Mtb* control. However, whether elevated IgM levels represented a biomarker or contributed directly to anti-microbial control remained unclear. Given the emerging data pointing to an anti-microbial role for polyclonal IgG and monoclonal IgG and IgA antibodies against *Mtb* (*32*, *39*–*44*), we next queried whether IgM also might harbor some anti-microbial capacity, using an engineered high-affinity LAM-specific antibody clone (A194) generated as an IgG1 and as an IgM (*45*).

In light of the previous observation that IgG1- and IgM-rich BAL from IV immunized rhesus macaques could drive intracellular *Mtb* killing in macrophages (Fig 3F), we first compared the anti-microbial activity of each isotype in a similar human monocyte-derived macrophage model. However, despite the anti-microbial signal observed in the BAL of IV BCG immunized animals, neither LAM-specific monoclonal antibody drove significant intracellular *Mtb* restriction in macrophages when added post-infection (Fig 6A).

**Figure 6:**
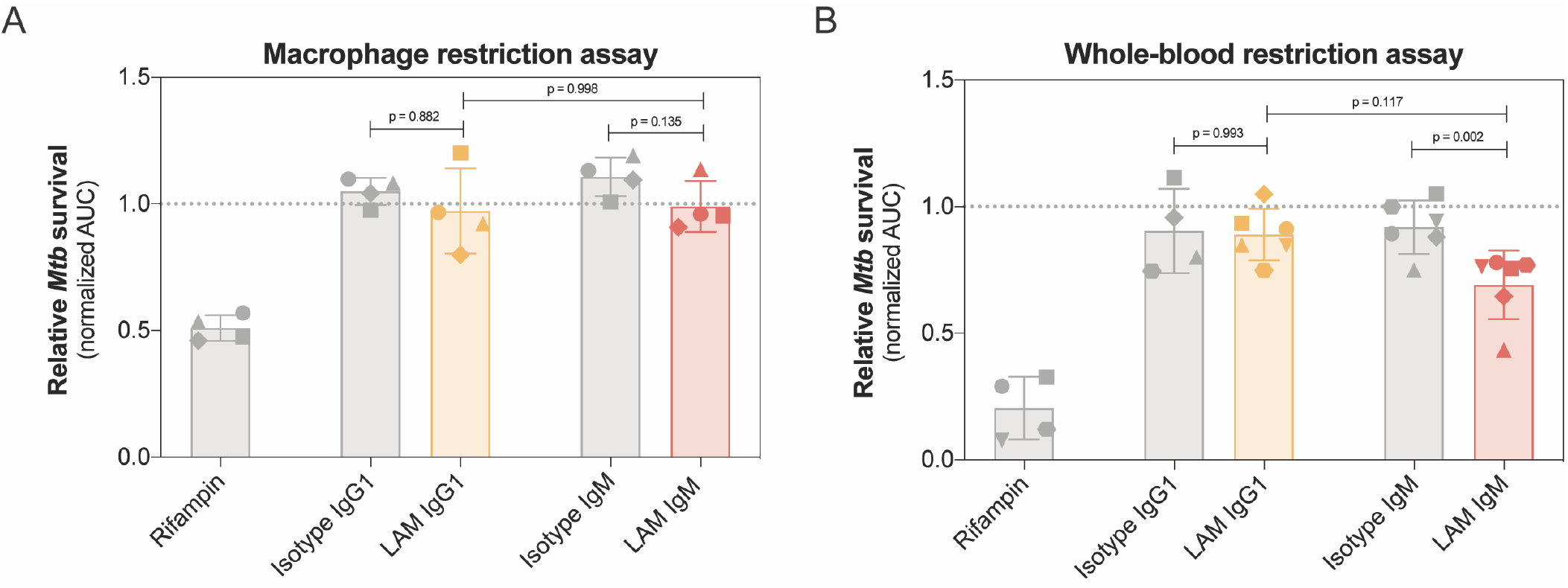
LAM IgM monoclonal antibody drives superior *Mtb* restriction in human whole-blood. (**A**) Macrophage restriction assay. Each antibody was added (final concentration 50ug/mL) to human monocyte-derived macrophages infected with a luminescent *Mtb* reporter strain (*Mtb-276*). Growth curves in the presence of each antibody treatment were generated by taking luminescence readings every 24 hours up to 120 hours. Y-axis is the area under the *Mtb* growth curve normalized by the no antibody condition of each donor. Each dot is the triplicate average from 1 donor. (**B**) Whole-blood restriction assay. Each antibody (final concentration 25ug/mL) was tested for their ability to drive *Mtb* restriction in the context of fresh human whole-blood using *Mtb-276*. Y-axis is the area under the *Mtb* growth curve, normalized by the no antibody condition. Growth curves were generated by taking luminescence readings every 24 hours up to 120 hours. Each dot is the triplicate average from 1 donor. Repeated measures ANOVA with Sidak’s multiple comparisons test.

While macrophages represent a primary cellular niche for *Mtb in vivo* during infection, we also probed the anti-microbial role for each LAM-specific antibody in a whole-blood model of infection – a system which queries the broader role of multiple immune cell types and components in microbial restriction. Specifically, fresh blood from healthy human donors was simultaneously infected with an *Mtb* luciferase reporter strain (*46*), and treated with each LAM-specific monoclonal antibody. Luminescence readings were then taken to obtain *Mtb* growth curves in the presence of each antibody treatment over the course of 120 hours. Remarkably, only the LAM-specific IgM antibody drove significant *Mtb* restriction in this system (Fig 6B). Further, the LAM-specific IgM antibody drove improved bacterial restriction compared to the IgG1 across nearly every donor tested (Fig 6B).

Ultimately, these data demonstrate that a high-affinity LAM-specific antibody clone drives improved *Mtb* restriction in whole-blood as an IgM, as compared an IgG1 variant, suggesting that in addition to representing an early marker of vaccine-induced *Mtb* control, *Mtb*-specific IgM antibodies have the potential to functionally contribute to immunologic control of *Mtb*.

## DISCUSSION

Recently, IV BCG vaccination in rhesus macaques was shown to result in robust protection against *Mtb* challenge (*20*), providing a unique opportunity to interrogate immunologic correlates and mechanisms of protection against *Mtb*. While published data highlighted the robust T cell immunity observed following IV BCG immunization, strong *Mtb* whole-cell lysate reactive humoral immune responses were also noted following this distinct vaccine delivery strategy (*20*). Given our emerging appreciation for a potential role for humoral immunity in *Mtb* control (*47*), here we deeply probed the antigen-specific humoral immune response across multiple BCG vaccine routes and doses to determine whether specific humoral immune profiles may complement cellular immunity, and potentially contribute to the protection afforded by IV BCG. Elevated antigen-specific antibody titers were observed in both the plasma and the lungs of IV BCG vaccinated animals, with a significant expansion of functional and anti-microbial responses in the BAL. Unexpectedly, correlation analyses revealed the unique association of BAL and plasma IgM responses close to or at the time of challenge, with reduced *Mtb* burden at necropsy in BCG immunized animals. Moreover, expanded plasma IgM titers were also observed in macaques immunized with an orthogonal, attenuated *Mtb* strain (*Mtb-ΔsigH*) that also conferred enhanced control over *Mtb* (*22*). Finally, a LAM-specific IgM antibody resulted in enhanced restrictive activity of *Mtb in vitro* compared to the same antibody clone with an IgG1 heavy chain, collectively pointing to a potential role for *Mtb*-specific IgM as a novel mechanistic correlate of protection in vaccine-induced *Mtb* control.

IV BCG immunized animals generated the strongest and most durable peripheral antibody responses directed to both protein antigens and to LAM. Conversely, standard ID BCG vaccination generated weaker antibody responses than the experimental vaccination regimens tested – each of which administered a larger dose of BCG (Fig 1). This pattern suggests that increased peak antibody titers may be a consequence of larger BCG dose delivered to these animals during immunization. However, each type of vaccination included in this study – intradermal, aerosol, and IV – additionally resulted in a distinct anatomic localization of vaccine antigen, where IV immunization resulted in robust localization in the spleen (*20*), a primary site of B cell activation (*48*). Thus, it is conceivable that an enrichment of antigen particularly in this primary B cell inductive site, may contribute to the strong and long-lasting peripheral humoral immunity uniquely observed in IV BCG immunized animals.

Peripheral antibody titers, often of the IgG isotype, represent a primary correlate of protection for the majority of approved vaccines (*47*, *49*–*55*). Yet surprisingly, vaccine-specific IgM titers were the only plasma antibody features to correlate inversely with *Mtb* burden in BCG immunized animals (Fig 4). Because protection was so dominantly associated with vaccination regimen, IgM titers did not represent an independent correlate of protection in the BCG route study. However, the potential value of IgM as a mechanistic correlate of immunity was corroborated by an enrichment in plasma IgM responses in animals that demonstrated enhanced *Mtb* control following *Mtb-ΔsigH* immunization (*22*). IgM plays a critical role during infection – particularly in the defense against other encapsulated bacteria – efficiently capable of driving phagocytosis, agglutination, and complement activation (*56*, *57*). IgM has also been recently implicated as a critical regulator of T cell immune responses (*58*), raising the prospect of both direct anti-microbial, and indirect cellular regulatory roles for IgM in immunity against *Mtb.*

In the present study, we observed that a high-affinity LAM-specific antibody exhibited superior anti-microbial activity in primary human whole-blood as an IgM compared to as an IgG1. Unlike IgM antibodies that are elicited by vaccination or infection, this IgM monoclonal was an engineered form of a relatively high affinity IgG antibody (A194-01) (*45*, *59*–*61*). While it is unclear whether IgM antibodies following IV BCG or *Mtb-ΔsigH* immunization require a similarly high affinity to drive anti-microbial function, the *in vitro* anti-microbial impact of this monoclonal suggests that IgM may not only serve as a surrogate of protective vaccine-induced immunity against *Mtb*, but may also play an unexpected mechanistic role in combating *Mtb* infection. Interestingly, several IgM LAM-specific monoclonal antibodies have been isolated from TB patients that also bind with moderate affinities, but not when expressed with an IgG heavy chain (*45*), suggesting that the increase in avidity provided by multimeric IgM may allow the antibody to access epitopes or drive Fc-mediated functions key to protective humoral immunity. Notwithstanding, future work should continue to dissect the mechanistic basis for this *Mtb* restriction activity – including Fab- and Fc-mediated mechanisms through which IgM may contribute to protection.

Beyond superior plasma antibody responses, IV BCG immunization was uniquely associated with a significant increase in BAL IgG1, IgA, and IgM antibody titers to all mycobacterial antigens tested at week 4 post-vaccination. Despite the highly functional nature of the BAL-derived antibodies following IV BCG vaccination, it remains unclear whether enhanced antibody titers and functions present in the BAL represent a signature of protection, a signature of IV vaccination, or both. Follow-up studies, using reduced dosing of IV BCG, may provide critical clues required to uncover the quantitative and qualitative correlates of immunity against *Mtb*. However, rhesus macaques vaccinated with repeated low-dose endobronchial instillation of BCG exhibited increased protection against *Mtb* challenge compared to macaques vaccinated with standard ID BCG (*62*). These protected animals exhibited significantly higher PPD-specific antibody titers in the BAL compared to animals that received intradermal BCG immunization. While only PPD-specific IgA and PPD-specific pan-isotype antibodies were measured, these results point again to robust lung-residing humoral immune responses as a common immune signature of protection between these two studies and across vaccination strategies. Notably, both intravenous and endobronchially instilled BCG have been shown to drive substantial BCG deposition in the lungs (*20*). As such, it is possible that the localization of vaccine antigen deep in lungs – rather than in the dermis by ID vaccination, or in the upper respiratory tract by AE vaccination – may be critical for the induction of robust, lung-specific T and B cell immunity, that may work together to interrupt infection. Lastly, despite the potent lung antibody responses observed in IV BCG immunized animals, the BAL antibody titers, functionality, and anti-microbial activity were largely transient (Figs 2 and 3). Thus, unlike lung T cell responses, which remained expanded over the course of the entire vaccination study (*20*), antibodies were less abundant in the lungs at the time of *Mtb* challenge (week 24). Critically, while differences may exist between BAL-derived antibodies and those found in the lung parenchyma, select antibody features – including LAM-specific IgG1, IgA, IgM titers, and LAM-specific FcγR3A binding antibodies – remained detectable and significantly higher in the airways of the IV group close to challenge. Given the small number of bacteria used in challenge (10 CFUs) (*20*), as well as the limited number of bacteria believed to cause infection in humans (*63*), it is plausible that even low levels of antibodies at the site of infection may be sufficient to capture the pathogen and contribute to first line defense. Of note, work in the context of influenza has demonstrated that vaccine-induced lung-resident memory B cell cells particularly IgM+ memory B cells – may also play a critical role in rapid response to infection, swiftly generating antibody-secreting cells that rapidly re-populate the lung with antibodies able to control and clear infection (*64*, *65*). Thus, it is conceivable that antibodies present in the airways 4 weeks post-vaccination, mark the establishment of lung-resident B cell immunity, which could respond instantaneously to *Mtb* challenge and contribute to protection in coordination with lung-resident T cell responses.

Ultimately, taken together, this work illustrates that IV BCG immunization drives superior plasma and lung antibody responses compared to ID and AE BCG vaccination. Specific humoral features associated with protection were identified across studies, highlighting potentially conserved roles for antibodies in vaccine-mediated protection against *Mtb*. Because of the potential safety issues associated with IV immunization, the development of alternative vaccination strategies able to mimic the protective humoral immune responses identified herein, may obviate the need for IV immunization to drive protection against TB in human populations. Thus, while efforts to leverage the immune response to combat TB via vaccination have largely focused on cellular immunity, this work demonstrates the value of a comprehensive examination of antibody characteristics across TB vaccine platforms, and motivates the continued study of antibodies as markers, and as functional mediators of protection against TB.

## MATERIALS AND METHODS

### Study design

Rhesus macaque (*Macaca mulatta*) plasma and bronchoalveolar lavage fluid (BAL) samples from the BCG route vaccination cohort were collected during a study performed at the Vaccine Research Center at the National Institutes of Health (*20*). All experimentation and sample collection from the original study complied with ethical regulations at the respective institutions (*20*). 48 BCG immunized animals were included in the study including: 10 animals that received standard intradermal (ID) BCG vaccination (target dose: 5 × 10^5^ CFUs), 8 animals that received high-dose intradermal (ID_high_) BCG vaccination (target dose: 5 × 10^7^ CFUs), 10 animals that received intravenous (IV) BCG vaccination (target dose: 5 × 10^7^ CFUs), 10 animals that received aerosol (AE) BCG vaccination (target dose: 5 × 10^7^ CFUs), and 10 animals that received a combination of AE and standard ID (AE/ID) BCG vaccination (target dose: AE 5 × 10^7^ CFUs, ID 5 × 10^5^ CFUs) (*20*). Following BCG vaccination, each macaque was challenged with 10 CFUs of *Mtb Erdman*, with a study endpoint of 12 weeks following *Mtb* challenge (*20*). In this study, *Mtb* burden values used throughout represent total thoracic CFUs measured at necropsy in the original study, and were measured as described previously (*20*). Plasma samples were analyzed from the following timepoints: pre-vaccination, week 8 post BCG vaccination, time of challenge (week 24 post BCG vaccination), and week 28 (4 weeks post *Mtb* challenge). BAL samples were analyzed from the following timepoints: pre-vaccination, week 4 post BCG vaccination, and week 16 post BCG vaccination. BAL was received as a 10X concentrate, and further diluted for experiments.

Rhesus macaque plasma samples from the attenuated *Mtb* vaccination cohort were collected during a study performed at the Tulane National Primate Research Center (*22*). All experimentation and sample collection from the original study were approved by the Institutional Animal Care and Use Committee and were performed in strict accordance with National Institutes of Health guidelines (*22*). Plasma from 9 rhesus macaques were analyzed in the present study. 4 animals received AE BCG vaccination (target dose: 1,000 CFUs), and 5 animals received AE *Mtb-ΔsigH* – an attenuated *Mtb* strain in the CDC1551 genetic background – vaccination (target dose: 1,000 CFUs). Eight weeks post-vaccination, each animal was challenged with a target dose of 1,000 CFUs of *Mtb CDC1551* (*22*). Plasma samples were analyzed from the following timepoints: pre-vaccination, week 7 post-vaccination, and necropsy (week 15).

### Antigens

To profile humoral immune responses, a panel of BCG/*Mtb*-antigens were used: purified protein derivative (PPD) (Statens Serum Institute), HspX (provided by T. Ottenhoff), LAM (BEI Resources, NR-14848), PstS1 (BEI Resources, NR-14859), and Apa (BEI Resources, NR-14862). Zaire ebolavirus glycoprotein (R&D Systems) was used as a negative control for the attenuated *Mtb* analysis.

### Non-human primate reagents

Mouse anti-rhesus IgG1 (clone 7H11) and IgA (clone 9B9) secondary antibodies were obtained from the National Institutes of Health Nonhuman Primate Reagent Resource supported by AI126683 and OD010976. Mouse anti-monkey IgM (clone 2C11-1-5) was acquired from Life Diagnostics. Soluble rhesus macaque FcγR2A and FcγR3A were acquired from the Duke Human Vaccine Institute Protein Production Facility.

### Antigen-specific antibody levels

Magnetic carboxylated fluorescent beads of distinct regions (Luminex Corp.) were first coupled to each protein antigen in a two-step carbodiimide reaction as described previously (*23*). LAM was modified by 4-(4,6-dimethoxy[1,3,5]triazin-2-yl)-4-methyl-morpholinium (DMTMM) and coupled to Luminex magnetic carboxylated fluorescent beads using protocols described previously (*66*, *67*).

Luminex using antigen-coupled beads to measure relative levels of antigen-specific antibodies was then performed as described previously (*68*), with minor modifications. A master mix of antigen-coupled beads was made at a concentration of 16.67 beads per μL per region in 0.1% bovine serum albumin (BSA)-PBS, and 750 beads per region per well (45uL) were added to a clear, flat-bottom 384 well plate (Greiner). 5μL of diluted sample was then added to the wells. Plasma from the BCG route vaccination study was diluted and run at 1:10 and 1:100. The 1:100 dilution was utilized for LAM IgG1, Apa IgG1, LAM IgA, and all IgM antigens. The 1:10 dilution was utilized for the remaining conditions. BAL from the BCG route vaccination study was diluted as follows: IgG1 1X, IgA 1X, IgM 0.1X. Plasma from the attenuated *Mtb* vaccination study was diluted as follows: IgG1 1:150, IgA 1:150, IgM 1:750. After adding the diluted samples, the plate was incubated shaking at 700 RPM overnight at 4°C. Next, the plate was washed 6 times and 45uL of mouse anti-rhesus IgG1, IgA, or IgM antibody at 0.65ug/mL was added, and incubated shaking at 700 RPM at room temperature (RT) for 1 hour. The plate was then washed 6 times and 45uL of Phycoerythrin (PE)-conjugated goat anti-mouse IgG was added (ThermoFisher, 31861) and incubated shaking at 700RPM at RT for 1 hour. The plate was then washed 6 times, and resuspended in Sheath Fluid (Luminex Corp.) in a final volume of 60uL. PE median fluorescence intensity (MFI) levels were then measured via the FlexMap 3D (Luminex Corp.) Data are represented as fold change over pre-vaccination levels. Samples were measured in duplicate.

### Antigen-specific Fcγ receptor binding

Rhesus macaque FcγRs were biotinylated as described previously (*68*). In brief, each FcγR was biotinylated using a BirA biotin-protein ligase bulk reaction kit (Avidity) according to the protocol of the manufacturer, and excess biotin was removed using 3 kD cutoff centrifugal filter units (Amicon).

Luminex using biotinylated rhesus macaque FcγRs and antigen-coupled beads to measure relative binding levels of antigen-specific antibodies to FcγRs was then performed as described previously (*68*), with minor modifications. A master mix of antigen-coupled beads was made at a concentration of 16.67 beads per μL per region in 0.1% BSA-PBS, and 750 beads per region per well (45uL) were added to a clear, flat-bottom 384 well plate (Greiner). 5μL of sample (Plasma: FcγR2A 1:10, FcγR3A 1:10; BAL: FcγR2A 1X, FcγR3A 1X) was added to the wells and incubated shaking at 700 RPM overnight at 4°C. After overnight incubation, streptavidin-PE (ProZyme) was added to each biotinylated FcγR in a 4:1 molar ratio and incubated rotating for 10 minutes at RT. 500μM biotin was then added at 1:100 relative to the total solution volume to quench the extra streptavidin-PE, and incubated rotating for 10 minutes at RT. After washing the assay plate 6 times, 40μL of each prepared detection FcγR (1μg/ml in 0.1% BSA-PBS) was added to the immune-complexed microspheres and incubated shaking at 700 RPM for 1 hour at RT. The plate was then washed 6 times, and resuspended in Sheath Fluid (Luminex Corp.) in a final volume of 60uL. PE MFI levels were then measured via the FlexMap 3D (Luminex Corp.). Data represented as fold change over pre-vaccination level. Samples were measured in duplicate.

### Antibody-dependent cellular phagocytosis (ADCP)

PPD ADCP (data not shown) was measured as described previously (*32*, *69*). LAM ADCP was measured as described previously (*32*, *69*) with minor changes. For every 100ug of LAM (dissolved in ddH_2_O), 10uL of 1M sodium acetate (NaOAc), and 2.2uL of 50mM sodium periodate (NaIO_4_) was added. This oxidation reaction proceeded for 45 – 60min on ice in the dark. 12uL of 0.8M NaIO_4_ was then added to block oxidation, and the solution was incubated for 5 min at RT in the dark. Next, the oxidized LAM was transferred to a new tube, and 10uL of 1M NaOAc and 22uL of 50mM hydrazide biotin (Sigma) were added. This biotinylation reaction proceeded for 2 hours at RT. Excess biotin was then removed using Amicon Ultra 0.5.L columns (3K, Millipore Sigma) according to the instructions of the manufacturer. Biotinylated LAM was then added to FITC-conjugated neutravidin beads (Invitrogen, 1.0μm) at a ratio of 1μg antigen: 4μL beads, and incubated for overnight at 4°C. Excess antigen was washed away. Antigen-coated beads were incubated with 10uL of sample (plasma 1:10, BAL 1:1) for 2 hr at 37 °C. THP-1 cells (5 × 10^4^ per well) were added and incubated at 37 °C for 16 hr. Bead uptake was measured in fixed cells using flow cytometry on a BD LSRII (BD Biosciences) and analyzed by FlowJo 10.3. Phagocytic scores were calculated as: ((%FITC positive cells) x (geometric mean fluorescence intensity of the FITC positive cells)) divided by 10,000. Data are represented as fold change over pre-vaccination levels. Samples were run in duplicate.

### Antibody-dependent neutrophil phagocytosis (ADNP)

ADNP was performed as described previously (*70*), with minor changes. LAM was biotinylated and coupled to fluorescent neutravidin beads (1.0μm, Invitrogen), incubated with serum, and washed as described above for ADCP. During the 2-hour bead and serum incubation, fresh peripheral blood collected from healthy donors in acid citrate dextrose (ACD) anti-coagulant tubes was added at a 1:9 ratio to ACK lysis buffer (150mM NH_4_Cl, 8610mM KHCO_3_, 0.1mM Na_2_-EDTA, pH 7.4) for 5 minutes at RT. After red blood cell lysis, the blood was centrifuged for 5 minutes at 1500 RPM. After centrifugation, supernatant was removed, and leukocytes were washed with 50ml of 4°C PBS, spun for 5 minutes at 1500 rpm and resuspended in R10 medium – (RPMI (Sigma), 10% fetal bovine serum (Sigma), 10mM HEPES (Corning), 2mM L-glutamine (Corning)) – at a final concentration of 2.5 × 10^5^ cells/mL. Leukocytes (5 × 10^4^ cells/well) were then added to the immune-complexed beads and incubated for 1 hour at 37°C 5% CO_2_. Following this incubation, the plates were spun for 5 minutes at 500 × g. After removing the supernatant, anti-human CD66b-Pacific Blue (BioLegend) was added to the leukocytes, and the cells were incubated for 20 minutes at RT. Following this incubation, the cells were washed with PBS and fixed. Bead uptake was measured in fixed cells using flow cytometry on a BD LSRII (BD Biosciences) and analyzed by FlowJo 10.3. Phagocytic scores were calculated in the CD66b positive cell population. Data represented as fold change over pre-vaccination level. Samples were run in duplicate.

### Antibody-dependent NK cell activation (ADNKA)

ADNKA was performed as described previously (*32*), with minor changes. ELISA plates (Thermo Fisher, NUNC MaxiSorp flat bottom) were coated with 150ng/well of LAM and incubated overnight at 4°C. The plates were then washed with PBS and blocked with 5% BSA-PBS for 2 hours. Next, the plates were washed with PBS, and 50uL of sample (plasma 1:10, BAL 1X) was added and incubated for 2 hours at 37°C. One day prior to adding the diluted sample, NK cells were isolated from healthy donors using the RosetteSep human NK cell enrichment cocktail (Stemcell) and Sepmate conical tubes (Stemcell) according to the instructions of the manufacturer. Following isolation, NK cells were incubated overnight at 1.5 × 10^6^ cells/mL in R10 media with 1ng/mL human recombinant IL-15 (Stemcell). After the 2 hour serum incubation, the assay plates were washed, and 50,000 primary human NK cells, together with 2.5uL PE-Cy5 anti-human CD107a (BD), 0.4uL Brefeldin A (5mg/ml, Sigma), and 10uL GolgiStop (BD) were added to each well of the assay plates. The plates were then incubated for 5 hours at 37°C. Following the incubation, the samples from each well were stained with 1uL each of: PE-Cy7 anti-human CD56, APC-Cy7 anti-human CD16, and Alexa Fluor 700 anti-human CD3 (all from BD). After a 20 minute incubation at RT to allow extracellular staining, the plate was washed with PBS, and the cells were fixed using Perm A and Perm B (Invitrogen). The Perm B solution additionally contained PE anti-human MIP-1β, and APC anti-human IFNγ (both from BD) to allow intracellular cytokine staining. After a final wash in PBS, the cells were resuspended in PBS and the fluorescence of each marker was measured on a BD LSR II flow cytometer (BD Biosciences) and analyzed by FlowJo 10.3. NK cells were defined as CD3 negative, CD16 positive, CD56 positive cells. Data are represented as fold change over pre-vaccination levels. The assay was performed in biological duplicate using NK cells from 2 different donors.

### Macrophage restriction assay (*Mtb-live/dead*)

*In vitro* macrophage *Mtb* survival was measured as described previously (*32*), with minor changes. CD14 positive cells were isolated from HIV negative donors using the EasySep CD14 Selection Kit II according to the instructions of the manufacturer (Stemcell). CD14 positive cells were matured for 7 days in R10 media without phenol in low adherent flasks (Corning). Monocyte-derived macrophages (MDMs) were plated 50,000 cells per well in glass bottom, 96-well plates (Greiner) 24 hours prior to infection. A reporter *Mtb* strain (*Mtb-live/dead*) with constitutive mCherry expression and inducible green fluorescent protein (GFP) expression (*33*), was cultured in log phase and filtered through a 5μm filter (Milliplex) prior to MDM infection at a multiplicity of infection of 1 for 14 hours at 37°C. Extracellular bacteria were washed off, and 200uL of pooled sample from each of the vaccination groups diluted in R10 without phenol (plasma 1:100, BAL 1X) was added. 3 days following infection, anhydrotetracycline (Sigma) (200 ng/ml) was added for 16 hours to induce GFP expression. 96 hours following infection, cells were fixed and stained with DAPI. Data were analyzed using the Columbus Image Data Storage and Analysis System. Bacterial survival was calculated as the ratio of live to total bacteria (the number of GFP+ pixels (live) divided by the number of mCherry+ pixels (total burden)) within macrophages in each well. Bacterial survival for each condition was normalized by bacterial survival in the no antibody condition. The assay was performed in technical triplicate using MDMs from 4 different donors.

### Macrophage restriction assay (*Mtb-276*)

CD14 positive cells were isolated from HIV negative donors using the EasySep CD14 Selection Kit II according to the instructions of the manufacturer (Stemcell). CD14 positive cells were matured for 7 days in R10 media without phenol in low adherent flasks (Corning). MDMs were plated 50,000 cells per well in sterile, white, flat-bottom 96-well plate plates (Greiner) 24 hours prior to infection. An auto-luminescent *Mtb* reporter strain (*Mtb-276*) (*46*), was cultured in log phase and filtered through a 5μm filter (Milliplex) prior to MDM infection at a multiplicity of infection of 1, for 14 hours at 37°C. Extracellular bacteria were washed off, and each antibody treatment was diluted to 50ug/mL in R10 without phenol, and 200uL of diluted antibody was added to each MDM-containing well. Control treatments: rifampin (Sigma) at 1ug/mL, human IgG1 isotype control (BE0297, BioXcell) at 50ug/mL, and human IgM isotype control (31146, Invitrogen) at 50ug/mL. Luminescence readings were then taken every 24 hours, up to 120 hours following infection to obtain *Mtb* growth curves in the presence of each antibody treatment. Area under the curve values were then computed for each antibody treatment in GraphPad Prism (version 8.4.0).

### Whole-blood restriction assay

Whole-blood from HIV negative human donors was collected fresh the day of the experiment in acid citrate dextrose tubes. *Mtb-276* previously cultured in 7H9 media at 37°C in log phase was washed once and resuspended in R10 media without phenol. Whole-blood was then infected with *Mtb-276* such that the final concentration was 1×10^6^ bacteria per mL of blood. Immediately after adding *Mtb-276* to blood, 150uL of blood and 150uL of antibody samples pre-diluted to 50ug/mL in R10 media are added together into a sterile, white, flat-bottom 96-well plate in triplicate (Greiner). Final concentration of experimental antibody treatments: 25ug/mL. Final concentration of control treatments: rifampin (Sigma) 0.25ug/mL, human IgG1 isotype control (BE0297, BioXcell) 25ug/mL, and human IgM isotype control (31146, Invitrogen) 25ug/mL. Samples in each well are mixed, then the first luminescence reading is taken on a plate reader (Tecan Spark 10M). The plate is then incubated at 37°C. Every 24 hours post-infection for 120 hours, the samples in each well are mixed, and luminescence readings are taken on a plate reader to obtain *Mtb* growth curves in the presence of different antibody treatments. *Mtb* restriction in whole-blood is calculated as the area under the curve for each condition. Area under the curve values were computed for each antibody treatment in GraphPad Prism (version 8.4.0).

### LAM-specific monoclonal antibody expression

A194 LAM-specific antibodies were generated as described previously (*45*). In brief, A194-IgG1 was generated by transfecting the A194-IGG1VH and IGVK plasmids into Expi293 cells. A194-IgM was generated by transfecting the A194-IGM1VH, IGVK, and joining (J) chain plasmids into Expi293 cells to generate multimeric IgM. Each antibody was purified by affinity chromatography. Protein A beads and protein L beads were used for the purification of IgG1 and IgM respectively. The antibodies were eluted using a low pH buffer, and characterized by SDS-PAGE for purity and size.

### Partial least squares discriminant analysis (PLS-DA)

A multivariate model to distinguish protected and susceptible macaques was generated using a combination of least absolute shrinkage and selection operator (LASSO)-based feature selection (*36*, *71*), and partial least squares discriminant analysis (PLS-DA) (*37*, *38*). Protected macaques were defined as those with an *Mtb* burden less than 1000 CFU/mL at time of necropsy. *Mtb* burden values used represent total thoracic CFUs measured at necropsy in the original study, and were measured as described previously (*20*).

For feature selection, the data were z-scored and 100 bootstrap datasets were generated. A LASSO model in which the optimal penalty term lambda was chosen via 5-fold cross-validation, was then fit on each bootstrap dataset, and coefficients from each iteration of LASSO regularization were stored. Using these coefficients, variable inclusion probabilities – defined as the proportion of bootstrap replications in which a coefficient estimate is non-zero – were computed for each antibody feature. LASSO regularization was implemented using the glmnet package (version 3.0-2) in R (version 3.6.2).

PLS-DA models across a grid of variable inclusion probability cutoffs were fit in a 5-fold cross-validation framework repeated 100 times. Model accuracy – defined as ((1 – balanced error rate) × 100) – was computed for each. The optimal model, which contained 3 antibody features, was found at a variable inclusion probability of 0.45. A graph of the first and second latent variable (LV) from the optimal PLS-DA model is included, as is a variable importance in the projection (VIP) plot, indicating the relative contribution of individual features to separation along the first LV. The significance of the model was assessed using a permutation test. Specifically, the group labels of the macaques were randomly permuted. PLS-DA models were then fit and evaluated for model accuracy in a 5-fold cross-validation framework repeated 100 times. The accuracy of the real model was compared with that of the permuted model using a Mann-Whitney U test. PLS-DA models were implemented using the mixOmics package (version 6.10.9) in R (version 3.6.2).

### Statistics

For the antibody titer (Fig 1 and 2), FcγR binding (Fig 3 and S1), and functional measurements (Fig 3), from the BCG dose vaccination cohort, Kruskal-Wallis with Dunn’s multiple-comparison tests were performed on the fold change values at each timepoint, comparing each vaccination group to the standard ID BCG group. For the macrophage *Mtb* restriction assay (Fig 3), a repeated measures ANOVA with Dunnett’s multiple comparisons test was performed for each vaccination group, comparing pre-vaccination restrictive activity with that of each post-vaccination timepoint. For antibody titers in the attenuated *Mtb* vaccination cohort (*Mtb-ΔsigH*) (Fig 5 and S2), Mann-Whitney U tests were performed on the fold change values at each timepoint, comparing aerosol *Mtb-ΔsigH* to the aerosol BCG group. For the LAM-specific monoclonal antibody *Mtb* restriction assays, a repeated measures ANOVA with Sidak’s multiple comparisons test was performed to make the relevant statistical comparisons. These statistics were performed in GraphPad Prism (version 8.4.0). Spearman correlations between *Mtb* burden and individual antibody features were computed in R (version 3.6.2) (Fig 4). Adjusted p-values (q-values) were calculated using the Benjamini-Hochberg procedure (*35*).

## Acknowledgements

We thank the Vaccine research center at the National Institutes of Health for providing the rhesus macaque plasma and BAL samples from the BCG route vaccination study. We thank the Tulane National Primate Research Center for providing the rhesus macaque plasma samples from the attenuated *Mtb* (*Mtb-ΔsigH*) vaccination study. We thank the Harvard Medical School, Laboratory for Systems Pharmacology for allowing the use of their automated microscope.

## Funding

Ragon Institute of MGH, MIT, and Harvard and the SAMANA Kay MGH Research Scholar Program. Bill and Melinda Gates Foundation: OPP1156795.

Defense Advanced Research Projects Agency: W911NF-19-2-0017.

National Institutes of Health: U54CA225088, U2CCA233262, U2CCA233280, AI150171-01, and Contract No. 75N93019C00071.

## Author Contributions

E.B.I. – Conceptualization, Methodology, Software, Validation, Formal Analysis, Investigation, Data Curation, Writing Original Draft, Review and Editing, Visualization, Funding Acquisition; A.O. – Validation, Investigation, Review and Editing; P.A.D. – Conceptualization, Resources, Data Curation, Review and Editing; S.S. – Investigation, Review and Editing; A.C. – Methodology, Resources, Review and Editing; W.L. – Methodology, Review and Editing; W.H. – Validation, Investigation, Resources; S.M. Conceptualization, Methodology, Resources, Review and Editing; D.K. – Conceptualization, Methodology, Resources, Data Curation, Review and Editing; H.P.G – Investigation, Resources, Data Curation; J.L.F. – Conceptualization, Methodology, Resources, Review and Editing, Supervision; M.R. – Conceptualization, Resources, Review and Editing, Supervision; R.A.S. – Conceptualization, Methodology, Resources, Review and Editing, Supervision; A.P. – Conceptualization, Methodology, Resources, Review and Editing, Supervision; S.F. – Conceptualization, Methodology, Resources, Review and Editing, Supervision, Project Administration, Funding Acquisition; G.A. – Conceptualization, Methodology, Resources, Review and Editing, Supervision, Project Administration, Funding Acquisition.

## Competing Interests

Galit Alter is a founder of SeromYx Systems, Inc.

## Data and materials availability

All data associated with this study are available in the main text or in the supplementary materials. Any additional materials data, and code will be made available to members of the scientific community in a timely fashion following a reasonable request.

## List of Supplementary Materials

Figure S1: Antigen-specific FcγR binding capacity of plasma and BAL antibodies

Figure S2: Plasma IgG1 and IgA titers from the attenuated *Mtb* (*Mtb-ΔsigH*) vaccination cohort

Data S1: Data_SystemsSerology

Code S1: Code_LASSO_PLSDA

