## Supplemental Materials for "Robust IgM responses following vaccination are associated with prevention of *Mycobacterium tuberculosis* infection in macaques"

### SUPPLEMENTARY FIGURES

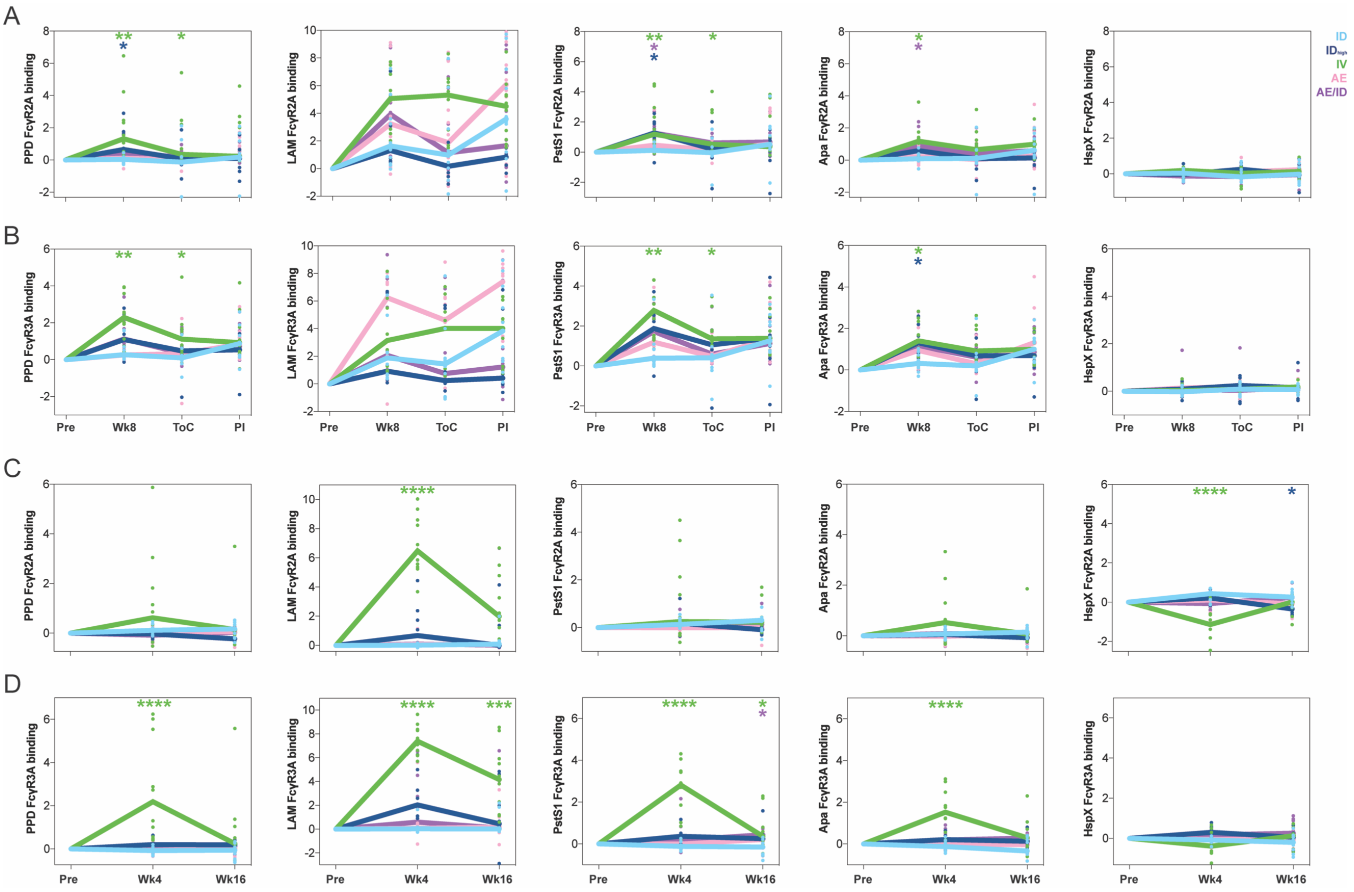


**Figure S1: Antigen-specific FcγR binding capacity of plasma and BAL antibodies.** Fold change in (**A and C**) FcγR2A, and (**B and D**) FcγR3A binding antibodies to PPD, LAM, PstS1, Apa, and HspX in the (**A and B**) plasma, and (**B and C**) BAL of each BCG-vaccinated rhesus macaque as measured via Luminex. Fold changes were calculated as fold change in Luminex median fluorescence intensity (MFI) over the pre-vaccination level for each primate. A base-2 log scale is used for the y-axis. Plasma timepoints: pre-vaccination (Pre), week 8 post-BCG vaccination (Wk8), time of challenge at week 24 post-BCG vaccination (ToC), post-infection at week 28 post-BCG vaccination (PI). BAL timepoints: pre-vaccination (Pre), week 4 post-BCG vaccination (Wk4), week 16 post-BCG vaccination (Wk16). Groups: standard intradermal BCG (light blue), high intradermal BCG (dark blue), intravenous BCG (green), aerosol BCG (pink), aerosol + intradermal BCG (purple). Each dot represents a single animal at the respective timepoint. The lines represent group medians over time. Kruskal-Wallis with Dunn's multiple-comparison tests were performed on the fold change values at each timepoint, comparing each vaccination group to the standard intradermal BCG group. Adjusted p-values are as follows: *, $p < 0.05$; **, $p < 0.01$; ***, $p < 0.001$; ****, $p < 0.0001$.

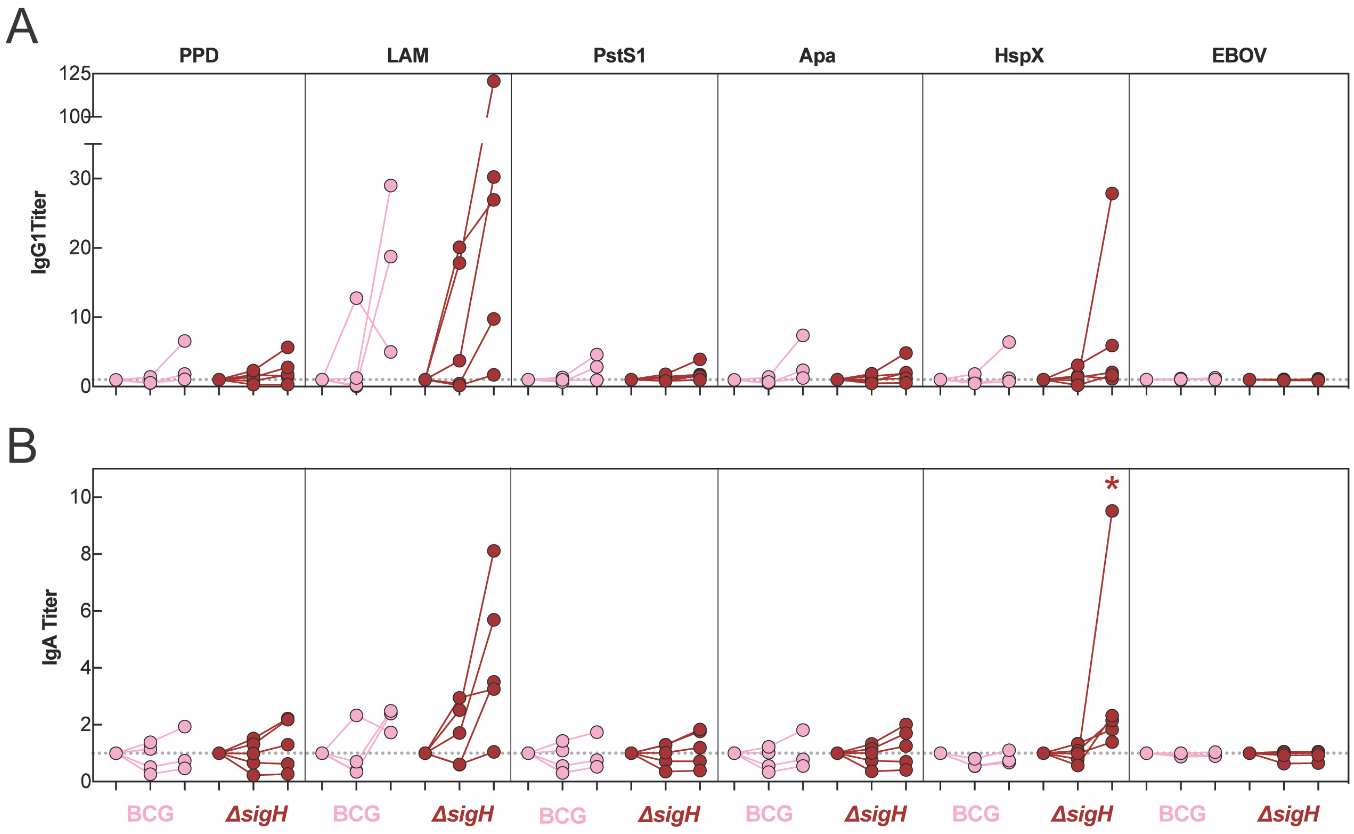


**Figure S2: Plasma IgG1 and IgA titers from the attenuated *Mtb* (*Mtb-ΔsigH*) vaccination cohort.** Fold change in (**A**) IgG1 and (**B**) IgA titers present in the plasma of each rhesus macaque following vaccination determined via Luminex. Fold changes were calculated as fold change in Luminex MFI over the pre-vaccination level for each primate. Each dot represents a single animal at the respective timepoint. Timepoints: pre-vaccination (left), week 7 post-vaccination (middle), week 15 post-vaccination at necropsy (right). *Mtb* challenge was performed week 8 post-vaccination. Mann-Whitney U test was performed on the fold change values at each timepoint, comparing aerosol *Mtb-ΔsigH* to the aerosol BCG group. p-values are as follows: *, $p < 0.05$; **, $p < 0.01$; ***, $p < 0.001$; ****, $p < 0.0001$.
